# Group A Streptococcus Infection of the Nasopharynx Requires Proinflammatory Signaling Through the Interleukin-1 Receptor

**DOI:** 10.1101/2020.06.12.149526

**Authors:** Doris L. LaRock, Raedeen Russell, Anders F. Johnson, Shyra Wilde, Christopher N. LaRock

## Abstract

Group A *Streptococcus* (GAS) is the etiologic agent of numerous high morbidity and high mortality diseases which commonly have a highly proinflammatory pathology. One factor contributing to this inflammation is the GAS protease SpeB, which directly activates the proinflammatory cytokine interleukin-1β (IL-1β), independent of the canonical inflammasome pathway. IL-1β drives neutrophil activation and recruitment that limits bacterial growth and invasion during invasive skin and soft tissue infections like necrotizing fasciitis. GAS also causes pharyngitis (strep throat), and the upper respiratory tract is its primary nidus for growth and transmission. Since the fitness selection for the species is likely primarily for this site, we examined the process of IL-1β activation in the murine nasopharynx. SpeB still activated IL-1β, which was required for neutrophil migration, but this inflammation instead increased GAS replication. Inhibiting IL-1β or depleting neutrophils, which both promote invasive infection, prevented GAS infection of the nasopharynx. Prior antibiotic exposure increased GAS growth in the murine nasopharynx, and antibiotics were sufficient to reverse the attenuation previously observed when IL-1β, neutrophils, or SpeB were not present to drive inflammation. Therefore, the same fundamental mechanism has opposing effects on virulence at different body sites. Invasive disease may be limited in part due to specific adaptations for inducing host inflammation that are beneficial for pharyngitis.

**IMPORTANCE:** Our previous reports showed that Group A *Streptococcus* (GAS) protease SpeB directly activates the host proinflammatory cytokine IL-1β and this restricts invasive skin infection. The upper respiratory tract is the primary site of GAS colonization and infection, but the host-pathogen interactions at this site are still largely unknown. We provide the first evidence that IL-1β-mediated inflammation promotes upper respiratory tract infection. This provides experimental evidence that the notable inflammation of strep throat, which presents with significant swelling, pain, and neutrophil influx, is not an ineffectual immune response, but rather is a GAS-directed remodeling of this niche for its pathogenic benefit.

## INTRODUCTION

Group A *Streptococcus* (GAS, *Streptococcus pyogenes*) is a top cause of infectious mortality and responsible for over half a million annual deaths (1). Death is primarily due to invasive diseases, including sepsis, necrotizing fasciitis, and toxic shock syndrome, or autoimmune diseases, most prominently acute rheumatic fever and rheumatic heart disease. The nasopharyngeal mucosa and associated lymphoid tissues is the most common site for infection (strep throat pharyngitis) and the primary carriage site for dissemination of GAS between individuals and to other sites of the body (2). Humans are transiently colonized by GAS throughout childhood but can be culture-positive at any point in their life, often without overt symptoms of disease (3, 4).

Infection starts at the mucosal and epidermal surfaces where GAS adheres to and invades skin keratinocytes and epithelial cells (5–8) to gain access to lymphoid tissues (9). This leads to inflammation of the mucosa, swelling of the tonsils, and formation of white patches of neutrophilic pus, characteristic of pharyngitis. Many molecular details of the host-pathogen interactions at this site remain unknown. Similar to how some enteric pathogens translocate across intestinal barriers, GAS can penetrate deeper into the mucosa through M cells and by disrupting cell junctions (2). It is unknown to what extent cell and tissue invasion is an essential feature of pharyngitis; however, the intracellular population of bacteria is protected from some antibiotics and may act as a reservoir for recurrent infection (4).

GAS inoculated intranasally into mice have a similar tropism as in humans and adhere, colonize, and invade the nasopharyngeal mucosa and nasal-associated lymphoid tissues (NALT). Despite structural differences between murine NALT and human tonsils (9), the inflammation, neutrophil infiltration, and pathology in mice resembles human disease and has been a useful model for examining host-pathogen interactions (6, 9–17). Many virulence factors important at other body sites are also essential at this site, including capsule (18), superantigens (14), SpyCEP (19), ScpA (15), and the regulator CovRS (13). M protein, a multifunctional virulence factor anchored on the cell surface that is the target for serological typing, is dispensable (6).

Some GAS virulence factors act to induce inflammation. During nasopharyngeal infection, the superantigen SpeA forces T cell antigen receptors (TCR) to engage the major histocompatibility complex (MHC) class II molecules of antigen-presenting cells in an antigen-independent manner (14). This induces excessive T cell activation which is highly proinflammatory and promotes nasopharyngeal infection (20). During invasive skin infections, the protease SpeB is also strongly proinflammatory and directly activates the proinflammatory cytokine IL-1β, which is inert until an inhibitory domain is proteolytically removed, bypassing its ordinary activation by host caspase-family proteases (21). IL-1β represses bacterial growth during invasive skin infections; neutrophil ablation or IL-1β neutralization enhances GAS growth in murine models of invasive infection and is a risk factor for invasive infections in humans (21). GAS can evade IL-1β-mediated restriction by inactivating SpeB through spontaneous mutation in the CovRS/CsrRS regulators (21, 22), a frequent observation of isolates from invasive diseases but not pharyngitis (23–27).

Here, we use a murine model of disease to examine SpeB-mediated activation of IL-1β within the nasopharynx. In contrast to our observations in models of invasive skin and soft tissue infection, activation of IL-1β is not restrictive and instead promotes infection. IL-1 signaling is required for neutrophil recruitment, which is also required for infection, potentially to overcome microbial interference. Together, these results have allowed us to examine how inflammation can promote pathogenesis and helped decipher how the cost-benefit of this strategy for GAS changes by host immune statue and infection site.

## MATERIALS AND METHODS

### Bacterial strains

GAS M1T1 5448, its isogenic *ΔspeB, Lactococcus lactis*, and the *speB* complementation vector pSpeB are previously described (7, 21, 28). GAS strains were statically grown at 37 °Cand 5% CO_2_ in Todd Hewitt broth (Difco), washed two times with phosphate-buffered saline (PBS, pH 7.4), and diluted to a multiplicity of infection (MOI) of 10 or 100 for *in vitro* infections.

### Experimental infection

7 to 8-week-old wildtype (Jackson labs), C57BL/6 mice of both sexes were used for experiments; *caspase-1/11*^-/-^ and *IL1R1*^-/-^ C57BL/6 have been previously described (21). Experiment treatments to these mice included intravenous delivery of 50 mg/kg Anakinra (Kineret; inhibits both human and murine IL-1R1 (21)) 4 h pre-infection, intraperitoneal delivery of 5000U penicillin G (Pfizer) 24 h pre-infection or as indicated, neutrophil depletion with 100 µg αLy6G/Ly6C mAb or isotype IgG (both BioXCell) 24 h pre-infection. Mouse groups were routinely inoculated with intranasally with 10^8^ CFU GAS slowly administered via micropipette in 10 μl PBS shared between nostrils and allowed to aspirate the inoculum via the normal breathing process. This volume resulted in no respiration into the lung, which can cause a necrotizing pneumonia and rapid death. At the indicated intervals the mice are euthanized, nasopharynx lavaged with 100 μl PBS and used for quantification of cytokines, cytometry, and/or CFU.

### *Cytometry* and *Cytokine Measurements*

Cytokine levels in cell-free tissue homogenate was quantified by multiple ELISA with the Mouse proinflammatory panel 1, following the manufacturer’s instructions (K15048G; Meso Scale Diagnostics). Cells treated with GolgiStop (BD), fixed with 2% paraformaldehyde with Fc block (BD Biosciences) before and after, and incubated with αCD3-BV605, αCD11c-BUV, αCD11b-PECF594, αGr-1/Ly6-APCH7, αCD40-BV650, αCD80-BUV737, αCD86-APCR700 (all Biolegend), to look at T cells, monocytes, macrophages, dendritic cells, and neutrophils. Depletion was confirmed by analyzing populations in blood obtained by cardiac puncture, with Ly6G, CD11b, and CD11c, as above. Flow cytometry was performed on a BD LSRII or LSRFortessa X-20 and analyzed using FlowJo 20 (TreeStar).

Transgenic IL-1R reporter cells were used to measure SpeB-matured IL-1β essentially as previously (21). These reporter cells were modified as previously described (29) to use luciferase as the readout, which signal limits interference from other bacterial and murine alkaline phosphatases. Luciferase activity was measured with Steady-Luc luciferin (Biotium) on a multimode plate reader (PerkinElmer).

### SpeB activity

Total SpeB activity was measured in individual isolates by hydrolysis of azocasein (Sigma) by methods previously described (21, 30). At least twelve isolates from each biological replicate were examined, and phenotypic conversion to the *covS*^-^ (SpeB^-^) phenotype is expressed as the percentage of isolates with a heritable loss of casein proteolysis.

### Neutrophil isolation and infection

Blood was collected from healthy male and female human volunteers under informed consent with approval by the Emory University Institutional Review Board. Neutrophils were isolated using PolyMorphPrep (Axis-Shield) as previously described (31). Neutrophil viability and concentration was assessed microscopically using 0.04% trypan blue then diluted to 10^5^ in PBS and infected with 10^6^ CFU. Aliquots were removed when indicated, incubated 2 min with 0.02% Triton-X100, and plated for CFU.

### Statistical analysis

Statistical analyses were performed using Prism 8 (GraphPad). Values are expressed as means ± standard error unless otherwise specified. Differences were determined using the Mann-Whitney U test (paired) or the Kruskal-Wallis test with Dunn’s post-analysis (multiple groups) unless otherwise specified.

## RESULTS

### Anakinra antagonizes GAS colonization of the nasopharynx

Anakinra inhibition of IL-1 signaling increases GAS burden during murine invasive infection (21). Since GAS usually resides in the upper respiratory tract, we used the established murine nasopharyngeal infection model (6, 9, 13, 20) to assess Anakinra’s effect at this site. Mice administered Anakinra intravenously then inoculated intranasally had significantly reduced GAS titers (**Figure 1A**). Using multiplexed enzyme-linked immunosorbent assay (ELISA), proinflammatory cytokines were quantified and IL-1β, IL-6, TNF-α, and IL-12 were all found to be significantly induced by infection (**Figure 1B**). Anakinra treatment reduced IL-1β and IL-6. These effects are likely due to both reduced bacterial burden and IL-1β-mediated induction of IL-6 and autoregulation (32), but other cytokines were not significantly impacted (**Figure 1B**). IL-1β retaining the amino-terminal inhibitory domain is detected by ELISA even though it lacks proinflammatory activity, so we quantified how much proinflammatory signal was present using transgenic IL-1 receptor reporter cells that detect processes, active IL-1β but not the non-inflammatory full-length protein (21, 29); IL-1 signaling was completely inhibited by Anakinra (**Figure 1C**). IL-1α, another agonist for the IL-1 receptor that is typically membrane-anchored (32), was not present at detectable levels.

**Figure 1.**
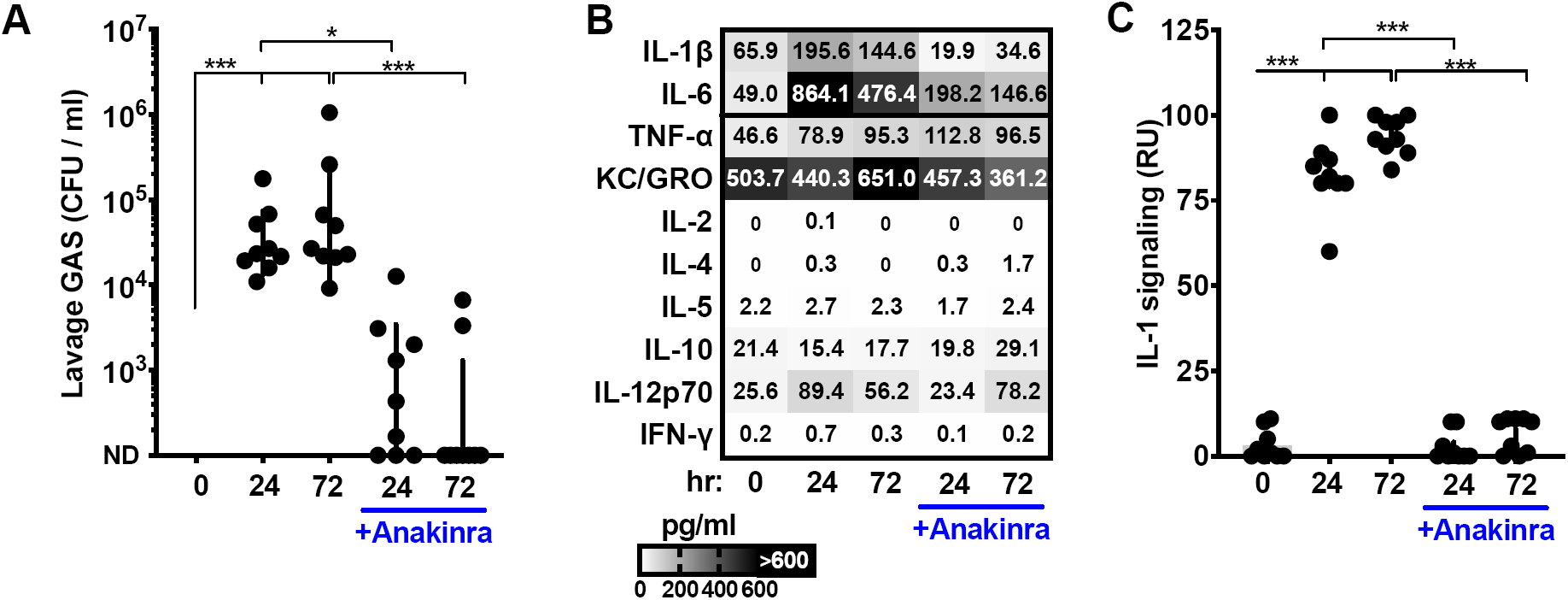
Anakinra antagonizes GAS colonization of the nasopharynx. Induction and inflammatory cytokine and their contribution to GAS infection. C57BL/6 mice treated with Anakinra (50 µg/kg) or PBS control and inoculated intranasally with 10^8^ colony forming units (CFU) of GAS M1T1 5448. Mice were euthanized after 24 or 72 h and the nasopharynx lavaged to (**A**) quantify GAS CFU by dilution plating, (**B**) quantify cytokines by MSD multiplex ELISA, and (**C**) measure levels of IL-1 signaling using a transgenic IL-1 receptor reporter specific for active IL-1α and IL-1β. Data are representative of at least 3 experiments, mean ± SD, N=10. ND, none detected; *, p<0.05; **, p<0.005; ***, p<0.0005.

### SpeB and Caspase-1 contribute to IL-1β generation in the nasopharynx

Similar to Anakinra treatment, mice deficient in the IL-1 receptor (*IL1R1*^-/-^) rapidly clear intranasally-inoculated GAS (**Figure 2A**). Levels of IL-1β were also reduced, consistent with both its autoregulation (32, 33) and the reduced burden of GAS. We further examined the pathways involved using mice lacking Caspase-1 and Caspase-11 (*casp-1/11*^-/-^), the inflammasome proteases that are typically essential for IL-1β maturation (32). Intranasally-inoculated Casp-1/11^-/-^ mice had a more modest, but significant, reduced GAS burden and induced less total IL-1β (**Figure 2A**). We have recently shown that in place of Caspase-1 and Caspase-11, the GAS protease SpeB can directly mature IL-1β and induce IL-1 signaling (21). Intranasally-inoculated Δ*speB* GAS induced significantly less total IL-1β, less active IL-1 signal, and was highly attenuated (**Figure 2B**). During skin infection, GAS growth and invasion is increased by Anakinra (21) but does not reverse the attenuation of Δ*speB* GAS in the nasopharynx (**Figure 2B**). Thus, inflammasome-associated Caspases also contribute to GAS growth in the nasopharynx, and not its restriction. However, this is not essential for GAS, unlike SpeB-mediated IL-1 signaling.

**Figure 2.**
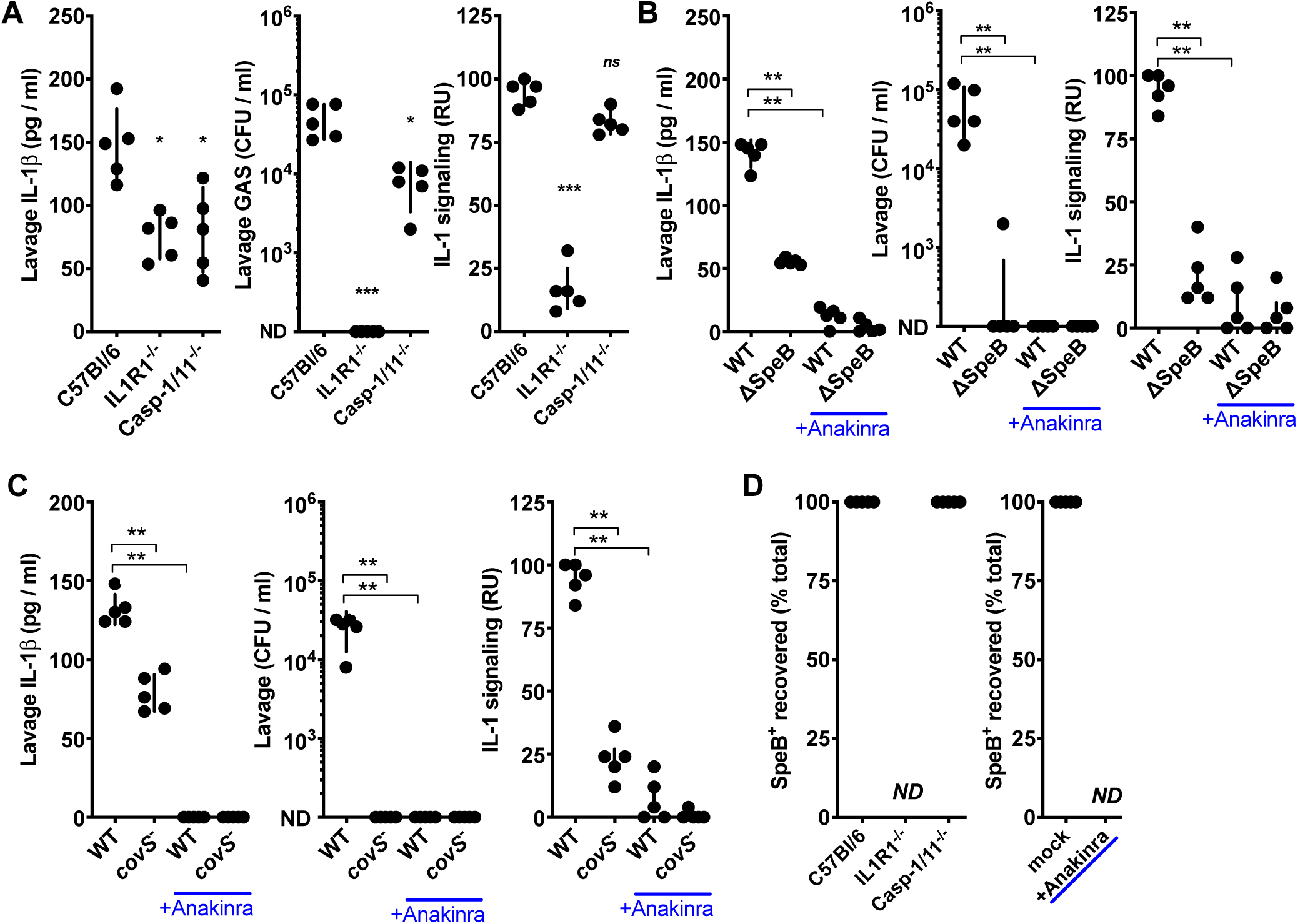
SpeB and Caspase-1 contribute to IL-1β generation in the nasopharynx. Effects of IL-1 and inflammasome signaling on GAS survival. (**A**) C57BL/6, *caspase-1/11*^-/-^, or *IL1R1*^-/-^ mice were inoculated intranasally with 10^8^ colony forming units wildtype GAS M1T1 5448. Mice were euthanized after 24 h and plated for CFU and cytokine levels quantified by ELISA and IL-1R reporter assay. (**B** and **C**) Role of SpeB and CovRS in GAS survival. C57BL/6 mice were administered Anakinra (50 µg/kg) or PBS, infected as above with wildtype, Δ*speB*, or AP *covS*^-^ GAS, and CFU and cytokines examined as above. (**D**) Role of host pathways in selection for *covS*^-^ SpeB^-^ clones. Isolated colonies from (**A, B**, and **C**) were screened for SpeB hydrolysis of azocaseine and the fraction where SpeB activity is lost indicated. ND, none detected, for experiments where no GAS were recoverable from the mouse. Data are representative of at least 3 experiments, mean ± SD, N=5. *, p<0.05; **, p<0.005; ***, p<0.0005.

To further investigate how SpeB contributes to infection of the nasopharynx we used an isogenic strain of M1T1 GAS (5448) developed a mutation inactivating CovS (*covS*^-^) during animal-passage (AP), as is frequently observed in isolates from human invasive infections (21, 25–27, 34). CovS mutation represses SpeB and greatly induces other important virulence factors including capsule, streptolysin O, and streptokinase, which during invasive infections may mitigate the attenuation due to loss of SpeB (21, 22). AP *covS*^-^ GAS induced less IL-1β and were attenuated in the nasopharynx, like Δ*speB* GAS (**Figure 2C**), and did not have the hypervirulent phenotype observed of *covS*^-^ GAS during invasive skin and soft-tissue infections (21, 26). *covS*^-^ (SpeB^-^) GAS spontaneously arise at high frequency during invasive infections, up to 10% of the recoverable bacteria at 24 h (21, 26, 35). No isolates of this genotype were detected in the nasopharynx and recovery was unaffected by IL-1 (*IL1R1*^-/-^ or Anakinra) or the inflammasome (*caspases-1/-11*^-/-^) (**Figure 2D**). Together these data show a requirement for SpeB and its essential regulator CovS in the nasopharynx that is not shared in the skin, and that this correlates with the ability to induce IL-1β.

### IL-1-dependent neutrophil recruitment promotes GAS nasopharyngeal infection

Since inflammation and GAS CFU were both decreased by Anakinra treatment, we examined which immune cells were altered by this treatment. After intravenous administration of Anakinra and intranasal inoculation with GAS, cells present at the infection site were removed by lavage and examined by flow cytometry. The only population significantly changed in frequency were Ly6G^+^CD11b^+^ neutrophils (**Figure 3A**), consistent with their robust recruitment during human infections (2). All neutrophils during GAS infection expressed high IL1R, which was lessened in Anakinra-treated mice (**Figure 3B**). While IL-1β may be driving neutrophil activation via this mechanisms, it is not a conventional chemokine, so it likely is promoting chemotaxis through the induction of IL-8 (CXCL8, murine homologs KC/MIP-2), CCL2, IFNγ, complement factors, and other known chemokines and their receptors (**Figure 1B**; (36, 37)).

**Figure 3.**
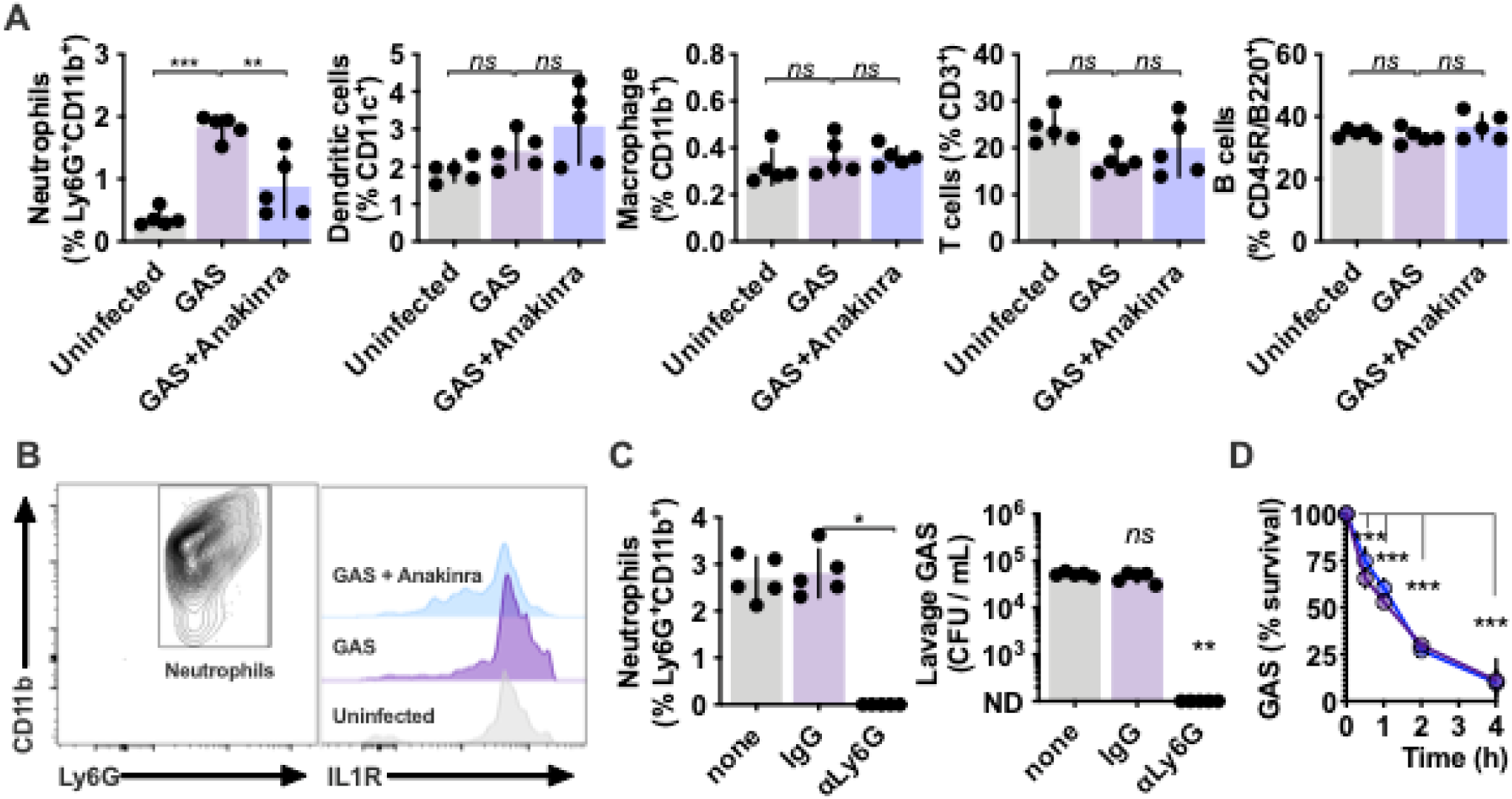
IL-1-dependent neutrophil recruitment promotes GAS nasopharyngeal infection. IL-1R signaling effects on nasopharyngeal cell populations. C57BL/6 mice treated with Anakinra (50 µg/kg) or PBS control inoculated intranasally with 10^8^ CFU of GAS M1T1 5448 and euthanized after 24 h (as in Figure 1A) and (**A**) nasopharyngeal lavage cells were analyzed by cytometry with the markers for neutrophils (CD11b^hi^Ly6G^hi^), dendritic cells (CD11c^hi^), macrophages (CD11b^hi^Ly6G^low^), T cells (CD3^+^), and B cells (CD45R^+^B220^+^) and expressed as a percentage of total live cells. (**B**) Effects of IL-1R signaling inhibition on IL-1R expression. Neutrophils (CD11b^hi^Ly6G^hi^) were further examined for expression of IL1R1 in each condition. (**C**) Neutrophil effects on GAS survival in the nasopharynx. Mice treated 24 h with αLy6G (100 µg), isotype IgG (100 µg), or PBS infected with 10^8^ CFU of GAS M1T1 5448 as above and neutrophils (CD11b^hi^Ly6G^hi^) and CFU quantified after 24 h. (**D**) Human neutrophils incubated with Anakinra (blue) or PBS only (purple) and infected with GAS at MOI 10. Aliquots were removed for CFU counts at the indicated intervals. Data are representative of at least 3 experiments, mean ± SD, N=5. ND, none detected; *, p<0.05; **, p<0.005; ***, p<0.0005.

Since neutrophil influx is a major feature of our model and natural infections, we next examined their contribution to disease by using an αLy6G antibody to deplete their numbers. While neutrophils are essential for restricting growth in models of invasive skin and soft tissue infection (21, 26, 38), neutrophil ablation completely blocked GAS infection of the nasopharynx (**Figure 3C**). Neutrophils effectively killed GAS *in vitro* (**Figure 3D**), suggesting that neutrophil recruitment may have compensatory benefits for GAS in the nasopharynx that do not occur in the skin and soft tissue.

### IL-1 mediates microbial interference in the nasopharynx

Nasopharyngeal infection with *Streptococcus pneumoniae* is inhibited by the anti-inflammatory corticosteroid dexamethasone (39). IL-1 signaling is not integral to this (40), as we find it is for GAS (**Figure 1A**), but we considered that inflammation more broadly may promote nasopharyngeal infection. The non-pathogenic Streptococcaceae species *Lactococcus lactis* was intranasally inoculated and after 24 h could be detected at similar levels in nasopharynx (**Figure 4A**). IL-1β was induced to significantly lessor levels by *L. lactis*, but was enhanced by *L. lactis* transformed to express SpeB (7) which led to clearance of the bacterium within 24 h (**Figure 4A**). The attenuation of SpeB-expressing *L. lactis* was reversed by Anakinra (**Figure 4B**), demonstrating that activation of IL-1β by SpeB is the major mechanism driving the attenuation of *L. lactis* + SpeB in the nasopharynx. Thus, inflammation that is beneficial to GAS is harmful to *L. lactis*. GAS and *L. lactis* intranasally coinocculated decreased recovery of both species (**Figure 4C**). During coculture in rich media (Todd-Hewitt broth), growth was not antagonistic (**Figure 4D**), indicating host factors are involved in the mutual antagonization between these species.

**Figure 4.**
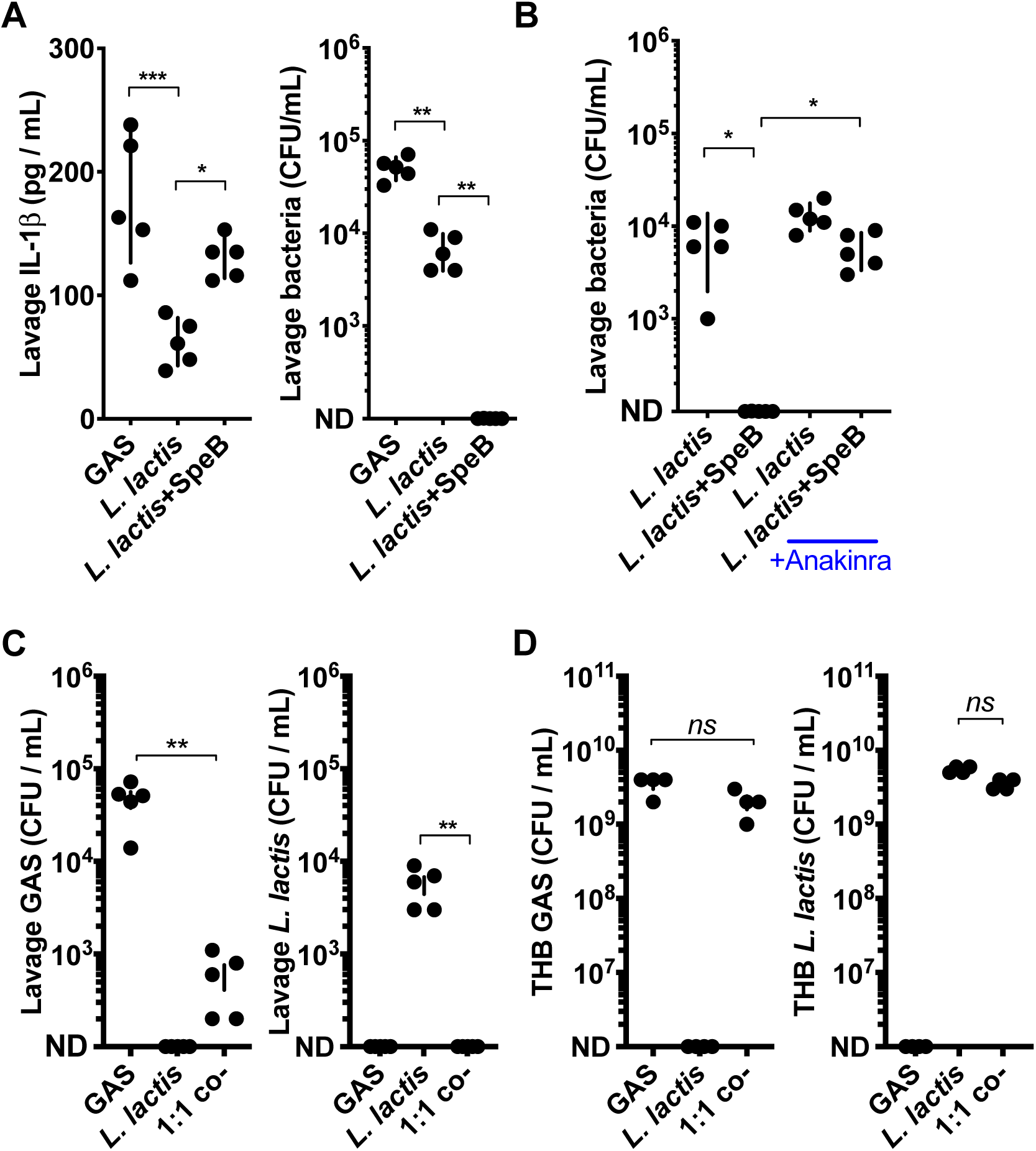
IL-1 mediates microbial interference in the nasopharynx. **(A)** SpeB-dependent effects on IL-1β and growth. C57BL/6 mice were inoculated intranasally with 10^8^ CFU of GAS, *L. lactis*, or SpeB-expressing *L. lactis*. IL-1β was quantified by ELISA and CFU by dilution plating from nasopharyngeal lavage samples collected after 24 h infection. **(B)** IL-1β-dependent effects on *L. lactis* growth. C57BL/6 mice treated with Anakinra (50 µg/kg) or PBS control were infected as above and CFU enumerated after 24 h as above. (**C**) *In vivo* competition experiment with GAS and *L. lactis*. C57BL/6 mice inoculated with 10^8^ CFU GAS, *L. lactis*, or a mix of both (5×10^7^ each; 1:1) and CFU enumerated after 24 h as above. (**D**) *In vitro* competition experiment with GAS and *L. lactis*. 10^4^ CFU GAS, *L. lactis*, or a mix of both (5×10^3^ each; 1:1) were grown 18 h in 3 mL THB at 37°C in 5% CO_2_, then CFU of each enumerated by dilution and differential plating. Data are representative of at least 3 experiments, mean ± SD, N=5. ND, none detected; *, p<0.05; **, p<0.005; ***, p<0.0005.

### Antibiotic pretreatment promotes GAS growth in the nasopharynx

The nasopharyngeal microbiota has to potential to interact with GAS in similar ways as *L. lactis*. Of note, early humans prospective studies show some microbiomes correlate with resistance to GAS (41) that is lost when their community structure is disrupted by penicillin treatment (42). Administration of penicillin as a single dose either 24 or 48 h before infection significantly increases GAS growth in the murine nasopharynx (**Figure 5A**), in agreement with these human studies. When the interval between penicillin treatment and infection is short, GAS does not survive due to its susceptibility. In mice pretreated with penicillin, the inhibition of GAS growth previously observed with Anakinra inhibition of IL-1 (**Figure 1A**) or neutrophil ablation (**Figure 4C**) is lost (**Figure 5B**). Furthermore, attenuation of Δ*speB* GAS in the nasopharynx (**Figure 2B**) is reversed in mice pre-treated with penicillin (**Figure 5C**). Thus, penicillin removes the proinflammatory requirements for SpeB, IL-1, and neutrophils during GAS infection of the nasopharynx.

**Figure 5.**
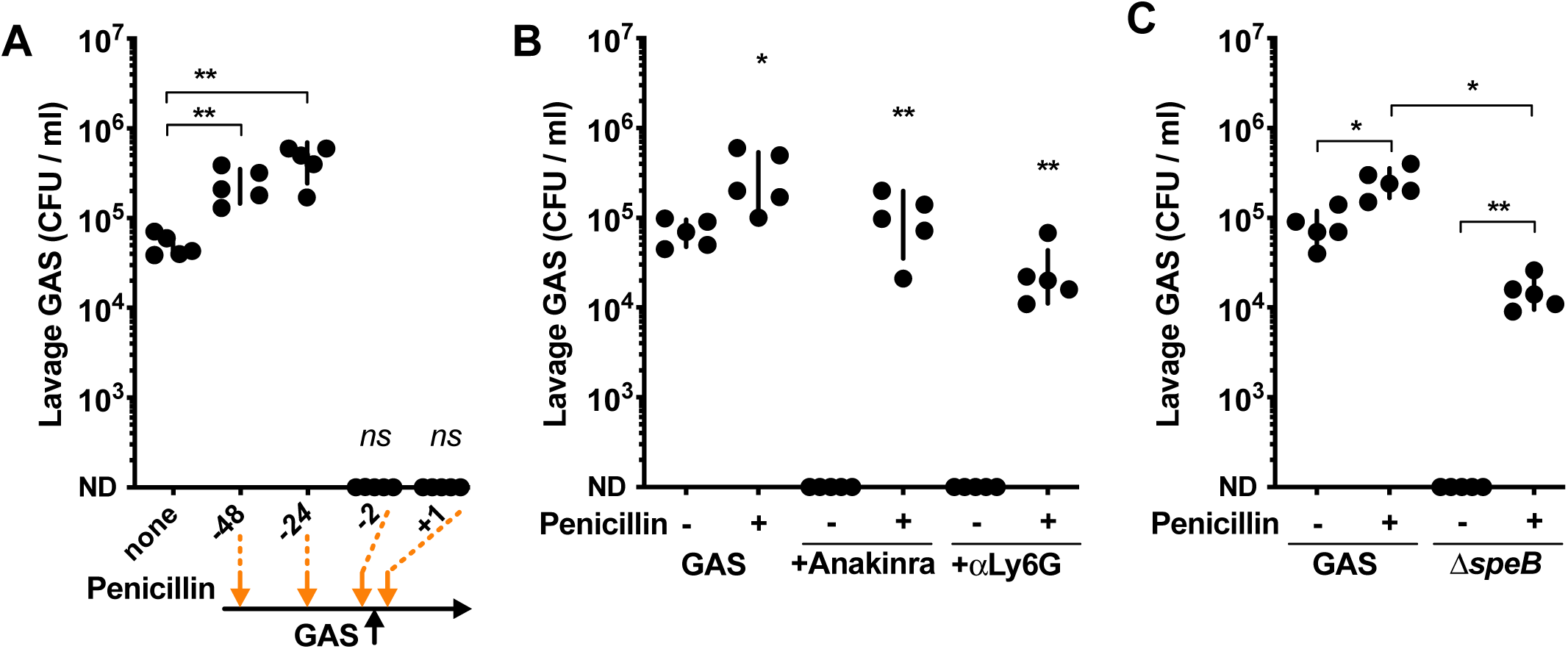
Antibiotic pretreatment promotes GAS growth in the nasopharynx. Growth of GAS in the upper respiratory tract of antibiotic pre-treated mice. (**A**) C57BL/6 mice were administered 5000U penicillin intraperitoneally at the indicated intervals (−48, -24, -2, or +1 h relative to inoculation time) and 10^8^ CFU of GAS 5448 delivered intranasally. Mice were euthanized 24 h post-inoculation and GAS CFU quantified from nasopharyngeal lavage. (**B**) C57BL/6 mice treated as above, with uniform administration of penicillin 24 h pre-infection, αLy6G 24 h pre-infection, or Anakinra 4 h pre-infection. (**C**) C57BL/6 mice were administered penicillin 24 h pre-infection as above, infected with 10^8^ CFU of wildtype or Δ*speB* GAS 5448 as above, and CFU enumerated after 24 h. Data are representative of at least 3 experiments, mean ± SD, N=5. *, p<0.05; **, p<0.005; ***, p<0.0005.

## DISCUSSION

IL-1 signaling induce antimicrobial effector mechanisms that restrict most pathogens, including GAS in invasive infections, and pharmacological inhibition of this critical proinflammatory pathway is a risk factor for severe infection in humans (21). We show that in the nasopharynx IL-1 signaling instead promotes GAS infection, highlighting a fundamental difference in inflammation and immunity between infection sites. Neutrophils are major immune cells recruited during infections of the nasopharynx and skin, and this is dependent on IL-1 signaling at both sites (21, 26, 32). Neutrophil influx in the skin and soft tissue is host-protective (2, 19, 38). However, in the nasopharynx, infection is promoted by this neutrophil infiltrate, which is a major component of strep throat pharyngitis.

Inflammation may be broadly beneficial for GAS in the nasopharynx. Other than SpeB, the superantigen SpeA also strongly induces inflammation by activating T cells; both the toxin and T cells are required for infection in this model (20, 43). Additional GAS proteins may similarly contribute to infection by not only acting as convention virulence factors, but also promoting inflammation as pathogen-associated molecular patterns (PAMPs. The strong resistance of GAS to immune effectors may be in part to survive this hyperinflammatory state induced by SpeB and superantigens, and decreased selection for immune invasion would occur in a less-inflamed host. *covS* frameshift mutations that repress *speB* and greatly induce most other virulence factors readily arise during invasive skin and soft tissue infections of human (23, 24) and mice (25). These mutations do not occur in human (34, 44, 45) or murine upper respiratory infections (21, 26, 27, 46, 47), supporting our observed requirement for CovS and SpeB in the nasopharynx. *covS* mutants also induce less IL-1β, less neutrophil infiltration, and are more resistant to neutrophils and other immune cells (21, 26, 48, 49). Accordingly, IL-1β and neutrophils are essential for *covS*^*-*^ selection during invasive skin infections (21, 48), though we report a reciprocal role in the nasopharynx. Since *covS* mutations are not fixed in the population, despite these seemingly beneficial activities that are advantageous at other sites, we infer that attenuation in the nasopharynx is a bottleneck exerting strong selection on the species.

Diverse other pathogens activate specific inflammatory pathways to antagonize competing microbes, disrupt membrane barrier function, promote dissemination, or acquire nutrients. Most notably, *S. pneumoniae* infection of the nasopharynx is inhibited by the broadly immunosuppressant corticosteroid dexamethasone (39), but not IL-1 signaling (50). IL-1 signaling still promotes neutrophil infiltration during *S. pneumoniae* infection, suggesting the underlying roles for inflammation in disease differ between these species. Other microbes of the upper respiratory tract including the pathogens *Haemophilus influenzae, Staphylococcus aureus, Moraxella catarrhalis*, and *Neisseria meningitidis*, and the microbiota, primarily other *Streptococcus, Haemophilus*, and *Neisseria* species (51), may similarly be impacted by inflammation and immunomodulation.

Consistent with our model that penicillin restores GAS growth, previous studies examining treatment failure with penicillin (52), recurrent infection (41), and infection secondary to antibiotics for other indications (42), support a role of antibiotics in human susceptibility. GAS colonization may be mediated by the nasopharyngeal microbiota. Resident species are able to directly kill GAS *in vitro* (53) and correlate with resistance to GAS infection (41) that is lost when their community structure is disrupted by penicillin treatment (42). Recapitulating this observation in the animal model we find penicillin increases susceptibility to GAS in the nasopharynx and removed the replication defects we observed upon targeted ablation of inflammatory responses. Together these results suggest that the distinctive inflammatory pathology of strep throat is key for pathogenesis. This establishes a connection between inflammation, antibiotics, and host permissiveness that has immediate clinical implications for this challenging pathogen.

## Acknowledgments

This work was supported by National Institute of Health NIH/NIAID K22 AI130223 (C.L.) and startup funds from Emory University. We thank Victor Nizet for bacterial strains, the Emory Multiplexed Immunoassay Core, the Children’s Healthcare of Atlanta and Emory University’s Children’s Clinical and Translational Discovery Core, and the Jacob Kohlmeier lab for technical support, and Jacqueline Kimmey and all members of the LaRock lab for critical manuscript review.

## REFERENCES

1. Carapetis JR, Steer AC, Mulholland EK, Weber M. 2005. The global burden of group A streptococcal diseases. Lancet Infect Dis 5:685–694.

2. Walker MJ, Barnett TC, McArthur JD, Cole J, Gillen CM, Henningham A, Sriprakash KS, Sanderson-Smith ML, Nizet V. 2014. Disease Manifestations and Pathogenic Mechanisms of Group A Streptococcus. Clin Microbiol Rev.

3. Holmes MC, Williams R. 1954. The distribution of carriers of Streptococcus pyogenes among 2,413 healthy children. J Hyg (Lond).

4. Kaplan EL, Chhatwal GS, Rohde M. 2006. Reduced Ability of Penicillin to Eradicate Ingested Group A Streptococci from Epithelial Cells: Clinical and Pathogenetic Implications. Clin Infect Dis Off Publ Infect Dis Soc Am 43:1398–1406.

5. Okada N, Liszewski MK, Atkinson JP, Caparon M. 1995. Membrane cofactor protein (CD46) is a keratinocyte receptor for the M protein of the group A streptococcus. Proc Natl Acad Sci U S A 92:2489–2493.

6. Anderson EL, Cole JN, Olson J, Ryba B, Ghosh P, Nizet V. 2014. The fibrinogen-binding M1 protein reduces pharyngeal cell adherence and colonization phenotypes of M1T1 group A Streptococcus. J Biol Chem 289:3539–3546.

7. Barnett TC, Liebl D, Seymour LM, Gillen CM, Lim JY, LaRock CN, Davies MR, Schulz BL, Nizet V, Teasdale RD, Walker MJ. 2013. The Globally Disseminated M1T1 Clone of Group A Streptococcus Evades Autophagy for Intracellular Replication. Cell Host Microbe.

8. Soderholm AT, Barnett TC, Korn O, Rivera-Hernandez T, Seymour LM, Schulz BL, Nizet V, Wells CA, Sweet MJ, Walker MJ. 2018. Group A Streptococcus M1T1 Intracellular Infection of Primary Tonsil Epithelial Cells Dampens Levels of Secreted IL-8 Through the Action of SpyCEP. Front Cell Infect Microbiol 8:160.

9. Park H-S, Francis KP, Yu J, Cleary PP. 2003. Membranous cells in nasal-associated lymphoid tissue: a portal of entry for the respiratory mucosal pathogen group A streptococcus. J Immunol 171:2532–2537.

10. Bessen D, Fischetti VA. 1988. Passive acquired mucosal immunity to group A streptococci by secretory immunoglobulin A. J Exp Med 167:1945–1950.

11. Dale JB, Baird RW, Courtney HS, Hasty DL, Bronze MS. 1994. Passive protection of mice against group A streptococcal pharyngeal infection by lipoteichoic acid. J Infect Dis 169:319–323.

12. Wessels MR, Bronze MS. 1994. Critical role of the group A streptococcal capsule in pharyngeal colonization and infection in mice. Proc Natl Acad Sci 91:12238–12242.

13. Alam FM, Turner CE, Smith K, Wiles S, Sriskandan S. 2013. Inactivation of the CovR/S Virulence Regulator Impairs Infection in an Improved Murine Model of Streptococcus pyogenes Naso-Pharyngeal Infection. PLoS ONE 8:e61655.

14. Kasper KJ, Zeppa JJ, Wakabayashi AT, Xu SX, Mazzuca DM, Welch I, Baroja ML, Kotb M, Cairns E, Cleary PP, Haeryfar SMM, McCormick JK. 2014. Bacterial Superantigens Promote Acute Nasopharyngeal Infection by Streptococcus pyogenes in a Human MHC Class II-Dependent Manner. PLoS Pathog 10:e1004155.

15. Park H-S, Cleary PP. 2005. Active and Passive Intranasal Immunizations with Streptococcal Surface Protein C5a Peptidase Prevent Infection of Murine Nasal Mucosa-Associated Lymphoid Tissue, a Functional Homologue of Human Tonsils. Infect Immun 73:7878–7886.

16. Dileepan T, Linehan JL, Moon JJ, Pepper M, Jenkins MK, Cleary PP. 2011. Robust Antigen Specific Th17 T Cell Response to Group A Streptococcus Is Dependent on IL-6 and Intranasal Route of Infection. PLoS Pathog 7:e1002252.

17. Wang B, Dileepan T, Briscoe S, Hyland KA, Kang J, Khoruts A, Cleary PP. 2010. Induction of TGF-beta1 and TGF-beta1-dependent predominant Th17 differentiation by group A streptococcal infection. Proc Natl Acad Sci U S A 107:5937–5942.

18. Vega LA, Sanson MA, Shah BJ, Flores AR. 2020. Strain-Dependent Effect of Capsule on Transmission and Persistence in an Infant Mouse Model of Group A *Streptococcus* Infection. Infect Immun 88:e00709-19, /iai/88/4/IAI.00709-19.atom.

19. Zinkernagel AS, Timmer AM, Pence MA, Locke JB, Buchanan JT, Turner CE, Mishalian I, Sriskandan S, Hanski E, Nizet V. 2008. The IL-8 Protease SpyCEP/ScpC of Group A Streptococcus Promotes Resistance to Neutrophil Killing. Cell Host Microbe.

20. Zeppa JJ, Kasper KJ, Mohorovic I, Mazzuca DM, Haeryfar SMM, McCormick JK. 2017. Nasopharyngeal infection by Streptococcus pyogenes requires superantigen-responsive Vβ-specific T cells. Proc Natl Acad Sci 114:10226–10231.

21. LaRock CN, Todd J, LaRock DL, Olson J, O’Donoghue AJ, Robertson AAB, Cooper MA, Hoffman HM, Nizet V. 2016. IL-1β is an innate immune sensor of microbial proteolysis. Sci Immunol 1:eaah3539–eaah3539.

22. Aziz RK, Pabst MJ, Jeng A, Kansal RG, Low DE, Nizet V, Kotb M. 2004. Invasive M1T1 group A Streptococcus undergoes a phase-shift in vivo to prevent proteolytic degradation of multiple virulence factors by SpeB. Mol Microbiol.

23. Chatellier S, Ihendyane N, Kansal RG, Khambaty F, Basma H, Norrby-Teglund A, Low DE, McGeer A, Kotb M. 2000. Genetic Relatedness and Superantigen Expression in Group A Streptococcus Serotype M1 Isolates from Patients with Severe and Nonsevere Invasive Diseases. Infect Immun 68:3523–3534.

24. Kansal RG, McGeer A, Low DE, Norrby-Teglund A, Kotb M. 2000. Inverse relation between disease severity and expression of the streptococcal cysteine protease, SpeB, among clonal M1T1 isolates recovered from invasive group A Streptococcal Infection Cases. Infect Immun.

25. Engleberg NC, Heath A, Miller A, Rivera C, DiRita VJ. 2001. Spontaneous mutations in the CsrRS two-component regulatory system of Streptococcus pyogenes result in enhanced virulence in a murine model of skin and soft tissue infection. J Infect Dis 183:1043–1054.

26. Walker MJ, Hollands A, Sanderson-Smith ML, Cole J, Kirk JK, Henningham A, McArthur JD, Dinkla K, Aziz RK, Kansal RG, Simpson AJ, Buchanan JT, Chhatwal GS, Kotb M, Nizet V. 2007. DNase Sda1 provides selection pressure for a switch to invasive group A streptococcal infection. Nat Med.

27. Cole J, Pence MA, von Köckritz-Blickwede M, Hollands A, Gallo RL, Walker MJ, Nizet V. 2010. M protein and hyaluronic acid capsule are essential for in vivo selection of covRS mutations characteristic of invasive serotype M1T1 group A Streptococcus. mBio 1.

28. LaRock CN, Döhrmann S, Todd J, Corriden R, Olson J, Johannssen T, Lepenies B, Gallo RL, Ghosh P, Nizet V. 2015. Group A Streptococcal M1 Protein Sequesters Cathelicidin to Evade Innate Immune Killing. Cell Host Microbe 1–9.

29. Sun J, LaRock DL, Skowronski EA, Kimmey JM, Olson J, Jiang Z, O’Donoghue AJ, Nizet V, LaRock CN. 2020. Role of Inflammasome-independent Activation of IL-1β by the Pseudomonas aeruginosa Protease LasB. bioRxiv 2020.05.18.101303.

30. Kwinn LA, Khosravi A, Aziz RK, Timmer AM, Doran KS, Kotb M, Nizet V. 2007. Genetic characterization and virulence role of the RALP3/LSA locus upstream of the streptolysin s operon in invasive M1T1 Group A Streptococcus. J Bacteriol 189:1322–1329.

31. Dohrmann S, LaRock CN, Anderson EL, Cole JN, Ryali B, Stewart C, Nonejuie P, Pogliano J, Corriden R, Ghosh P, Nizet V. 2017. Group A Streptococcal M1 Protein Provides Resistance against the Antimicrobial Activity of Histones. Sci Rep 7:43039.

32. LaRock CN, Nizet V. 2015. Inflammasome/IL-1 β responses to streptococcal pathogens. Front Immunol 1–12.

33. Toda Y, Tsukada J, Misago M, Kominato Y, Auron PE, Tanaka Y. 2002. Autocrine induction of the human pro-IL-1 beta gene promoter by IL-1 beta in monocytes. J Immunol 168:1984–1991.

34. Sumby P, Whitney AR, Graviss EA, DeLeo FR, Musser JM. 2006. Genome-wide analysis of group a streptococci reveals a mutation that modulates global phenotype and disease specificity. PLoS Pathog 2:e5.

35. Maamary PG, Zakour NLB, Cole J, Hollands A, Aziz RK, Barnett TC, Cork AJ, Henningham A, Sanderson-Smith ML, McArthur JD, Venturini C, Gillen CM, Kirk JK, Johnson DR, Taylor WL, Kaplan EL, Kotb M, Nizet V, Beatson SA, Walker MJ. 2012. Tracing the evolutionary history of the pandemic group A streptococcal M1T1 clone. FASEB J.

36. LaRock CN, Cookson BT. 2013. Burning Down the House: Cellular Actions during Pyroptosis. PLoS Pathog 9:e1003793.

37. Dinarello CA, Simon AK, van der meer JWM. 2012. Treating inflammation by blocking interleukin-1 in a broad spectrum of diseases. Nat Rev Drug Discov 11:633–652.

38. Hidalgo-Grass C, Mishalian I, Dan-Goor M, Belotserkovsky I, Eran Y, Nizet V, Peled A, Hanski E. 2006. A streptococcal protease that degrades CXC chemokines and impairs bacterial clearance from infected tissues. EMBO J 25:4628–4637.

39. Zafar MA, Wang Y, Hamaguchi S, Weiser JN. 2017. Host-to-Host Transmission of Streptococcus pneumoniae Is Driven by Its Inflammatory Toxin, Pneumolysin. Cell Host Microbe 21:73–83.

40. Lemon JK, Weiser JN. 2015. Degradation products of the extracellular pathogen Streptococcus pneumoniae access the cytosol via its pore-forming toxin. mBio 6.

41. Crowe CC, Sanders WE. 1973. Bacterial interference. II. Role of the normal throat flora in prevention of colonization by group A Streptococcus. J Infect Dis.

42. Sanders CC, Sanders WE, Harrowe DJ. 1976. Bacterial interference: effects of oral antibiotics on the normal throat flora and its ability to interfere with group A streptococci. Infect Immun 13:808–812.

43. Dan JM, Havenar-Daughton C, Kendric K, Al-kolla R, Kaushik K, Rosales SL, Anderson EL, LaRock CN, Vijayanand P, Seumois G, Layfield D, Cutress RI, Ottensmeier CH, Arlehamn CSL, Sette A, Nizet V, Bothwell M, Brigger M, Crotty S. 2019. Recurrent group A Streptococcus tonsillitis is an immunosusceptibility disease involving antibody deficiency and aberrant TFH cells. Sci Transl Med 11:eaau3776.

44. Ikebe T, Ato M, Matsumura T, Hasegawa H, Sata T, Kobayashi K, Watanabe H. 2010. Highly Frequent Mutations in Negative Regulators of Multiple Virulence Genes in Group A Streptococcal Toxic Shock Syndrome Isolates. PLOS Pathog 6:e1000832.

45. Shea PR, Beres SB, Flores AR, Ewbank AL, Gonzalez-Lugo JH, Martagon-Rosado AJ, Martinez-Gutierrez JC, Rehman HA, Serrano-Gonzalez M, Fittipaldi N, Ayers SD, Webb P, Willey BM, Low DE, Musser JM. 2011. Distinct signatures of diversifying selection revealed by genome analysis of respiratory tract and invasive bacterial populations. Proc Natl Acad Sci 108:5039–5044.

46. Fiebig A, Loof TG, Babbar A, Itzek A, Koehorst JJ, Schaap PJ, Nitsche-Schmitz DP. 2015. Comparative Genomics of Streptococcus pyogenes M1 isolates differing in virulence and propensity to cause systemic infection in mice. Int J Med Microbiol IJMM.

47. Horstmann N, Tran CN, Brumlow C, DebRoy S, Yao H, Nogueras Gonzalez G, Makthal N, Kumaraswami M, Shelburne SA. 2018. Phosphatase activity of the control of virulence sensor kinase CovS is critical for the pathogenesis of group A streptococcus. PLOS Pathog 14:e1007354.

48. Li J, Liu G, Feng W, Zhou Y, Liu M, Wiley JA, Lei B. 2014. Neutrophils Select Hypervirulent CovRS Mutants of M1T1 Group A Streptococcus During Subcutaneous Infection of Mice. Infect Immun.

49. LaRock CN, Nizet V. 2015. Cationic Antimicrobial Peptide Resistance Mechanisms of Streptococcal Pathogens. Biochim Biophys Acta.

50. Lysenko ES, Ratner AJ, Nelson AL, Weiser JN. 2005. The Role of Innate Immune Responses in the Outcome of Interspecies Competition for Colonization of Mucosal Surfaces. PLoS Pathog 1:e1.

51. Lemon KP, Klepac-Ceraj V, Schiffer HK, Brodie EL, Lynch SV, Kolter R. 2010. Comparative analyses of the bacterial microbiota of the human nostril and oropharynx. mBio 1:1–9.

52. Eagle H. 1952. Experimental approach to the problem of treatment failure with penicillin: I. Group A streptococcal infection in mice. Am J Med.

53. Sanders E. 1969. Bacterial interference. I. Its occurrence among the respiratory tract flora and characterization of inhibition of group A streptococci by viridans streptococci. J Infect Dis 120:698–707.

